# “TRIF is a key inflammatory mediator of acute sickness behavior and cancer cachexia”

**DOI:** 10.1101/232280

**Authors:** Kevin G. Burfeind, Xinxia Zhu, Peter R. Levasseur, Katherine A. Michaelis, Mason A. Norgard, Daniel L. Marks

## Abstract

Hypothalamic inflammation is a key component of acute sickness behavior and cachexia, yet mechanisms of inflammatory signaling in the central nervous system remain unclear. We assessed the role of TRIF signaling in acute inflammation (lipopolysaccharide (LPS) challenge) and in a chronic inflammatory state (cancer cachexia). TRIFKO mice resisted anorexia and weight loss after peripheral (intraperitoneal, IP) or central (intracerebroventricular, ICV) LPS challenge and in a model of pancreatic cancer cachexia. Compared to WT mice, TRIFKO mice showed attenuated upregulation of *Il6, Ccl2, Ccl5, Cxcl1, Cxcl2,* and *Cxcl10* in the hypothalamus after IP LPS treatment, as well as attenuated microglial activation and neutrophil infiltration into the brain after ICV LPS treatment. Our results show that TRIF is an important inflammatory signaling mediator of sickness behavior and cachexia and presents a novel therapeutic target for these conditions.

## Introduction

Innate immune activation in response to various pathogens leads to systemic inflammation, inducing a distinct metabolic and behavioral paradigm that includes fever, weight loss, anorexia, and fatigue. This constellation of signs and symptoms, referred to as “sickness behavior” (Dantzer et al., 1998), is critical for combating infection and allows resources to be diverted to the immune system to fight pathogens. However, if sickness behavior is maintained in conditions of chronic inflammation, it can become maladaptive and manifest as cachexia. Cachexia is a devastating syndrome characterized by anorexia, increased catabolism of lean body mass, and lethargy (Argiles et al., 2010; Evans et al., 2008; Fearon et al., 2011). It is prevalent in numerous chronic diseases, including cancer (Tisdale, 2002), chronic renal failure (A. Y. Wang et al., 2004), congestive heart failure (Anker et al., 1997), and untreated HIV (Kotler, Tierney, Wang, & Pierson, 1989). Furthermore, cachexia is associated with increased mortality of the underlying disease and decreased quality of life (Bachmann et al., 2008; Lainscak, Podbregar, & Anker, 2007; Wesseltoft-Rao et al., 2015). Despite this serious clinical concern, there are currently no effective treatments and mechanisms remain controversial.

Our lab, along with others, described a central nervous system (CNS)-based mechanism of cachexia in which cytokines generated in the periphery are amplified and modified within the hypothalamus, leading to aberrant activity of weight- and activity-modulating neurons (Bluthé, Michaud, Poli, & Dantzer, 2000; Braun et al., 2011; Burfeind, Michaelis, & Marks, 2015; Grossberg et al., 2011). Specifically, intracerebroventricular (ICV) injection of inflammatory cytokines (Bodnar et al., 1989; Sonti, Ilyin, & Plata-Salaman, 1996) or pathogen associated molecular patterns such as lipopolysaccharide (LPS) (Wisse et al., 2007) potently reduces food intake and activity. Furthermore, peripheral or central cytokine injection or immune challenge leads to rapid activation of neurons in areas that are critical for food intake and energy metabolism, such as the nuclei of the mediobasal hypothalamus (MBH) (Elmquist, Scammell, Jacobsen, & Saper, 1996; Konsman, Tridon, & Dantzer, 2000; Laflamme & Rivest, 1999; Morgan & Curran, 1986). However, the cellular and molecular pathways whereby peripheral inflammation is translated in the brain into behavioral or metabolic responses remain undefined.

Toll-like receptors (TLRs) are key components of the innate immune system, recognizing a variety of pathogens and inflammatory signals. TLR function is important for mounting an appropriate inflammatory response, and metabolic signaling in the CNS is closely tied to TLR signaling (Jin et al., 2016b). Pro-inflammatory signaling via the Myeloid Differentiation Primary Response Gene 88 (MyD88) pathway was initially thought to be the dominant mechanism whereby the binding of pathogenic signaling molecules to receptors is linked to the synthesis and release of inflammatory cytokines and chemokines (Medzhitov et al., 1998). However, recent data suggest that MyD88-independent pathways linking TLRs to cellular activation are present within the brain (Hanke & Kielian, 2011; Sen Lin et al., 2012). The adaptor protein TIR-domain-containing adaptor inducing interferon-β (TRIF) is an important inflammatory signaling mediator, yet has received little attention in the context of CNS-mediated alterations in behavior and metabolism during illness. TRIF is the dominant adapter for TLR3 signaling, and plays an essential role in TLR4 responses to LPS as well (Yamamoto et al., 2003). Furthermore, TRIF knockout (TRIFKO) mice are nearly as resistant to endotoxin-induced mortality as are MyD88KO mice (Feng et al., 2011).

The role of TRIF signaling in the CNS during acute sickness behavior and cachexia is unknown. We found that TRIF signaling is important for neuroinflammation and resulting acute sickness behavior after systemic or central exposure to LPS. We also found that mice lacking TRIF have attenuated cancer cachexia. These results implicate TRIF as a key signaling mediator in inflammation-driven behavioral and metabolic changes during illness, and a potential therapeutic target for cachexia.

## Results

### Mice lacking TRIF show attenuated acute illness response after systemic LPS challenge

TRIF is an important adaptor protein for innate immune activation (Yamamoto et al., 2003). While several studies demonstrate that MyD88 is important for acute sickness behavior, the role of TRIF in sickness behavior after LPS challenge is unknown. After systemic LPS challenge (250 μg/kg, IP), TRIFKO mice showed attenuated anorexia and weight loss compared to WT mice (Fig. 1a and b). Next, in order to determine the degree of hypothalamic activation and quantify stress response, we measured plasma corticosterone (Gong et al., 2015). While WT mice showed a large increase in plasma corticosterone 4 hrs after IP LPS administration, LPS-treated TRIFKO mice did not show a significant increase (Fig. 1c).

**Figure 1:**
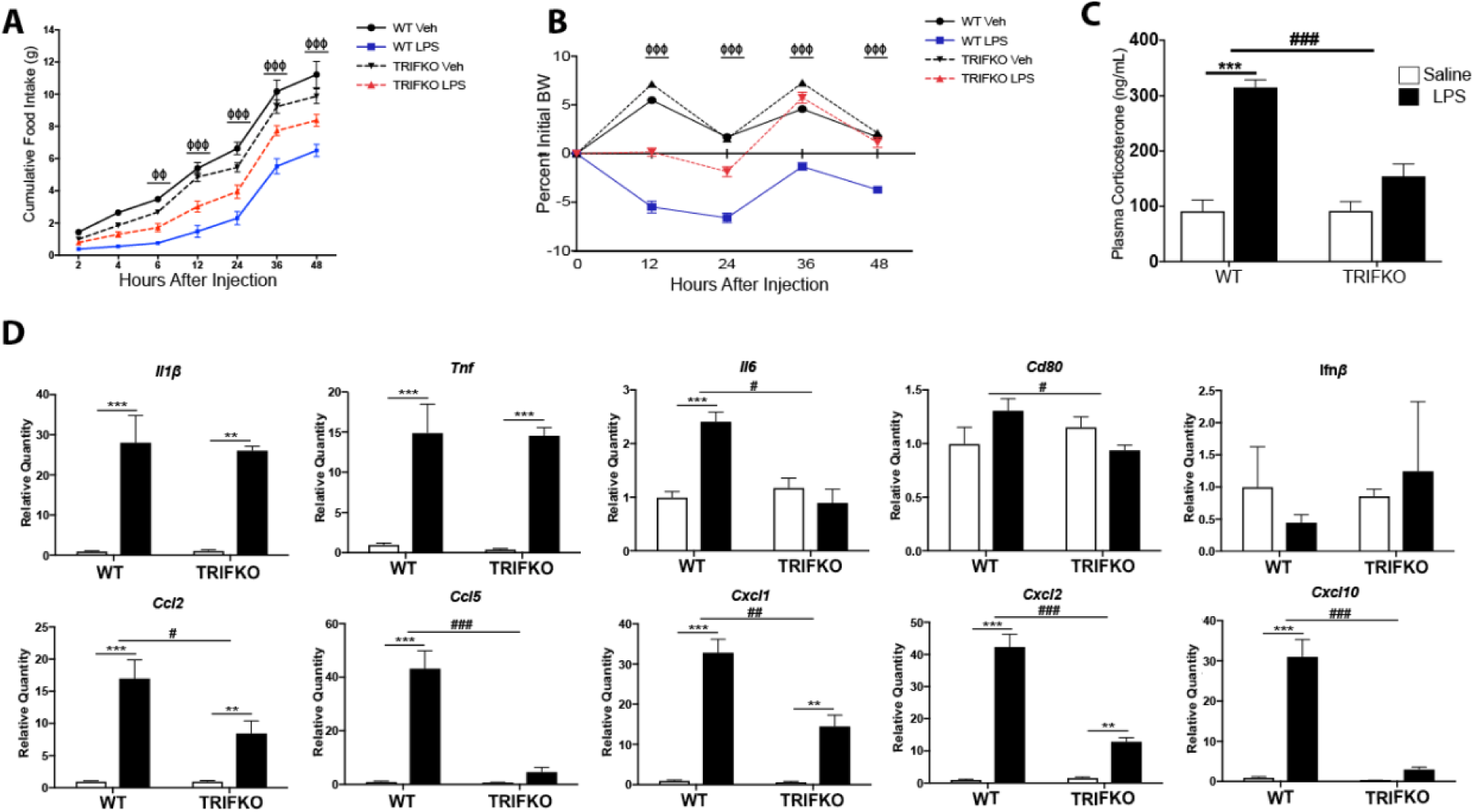
TRIFKO mice have attenuated acute sickness behavior in response to systemic LPS exposure. A) Cumulative food intake after 250 μg/kg IP LPS treatment. Veh = vehicle treatment (BSA/saline). Φ = p <.05, ΦΦ = p<.01, ΦΦΦ = p<.001 for WT LPS vs. TRIFKO LPS in Bonferroni post-hoc comparisons. B) Body weight change after 250 μg/kg IP LPS treatment. BW = body weight. ΦΦΦ = p<0.001 for WT LPS vs. TRIFKO LPS in Bonferroni post-hoc comparisons (See Figure 1 - source data 1 for food intake and body weight data). N = 5-7/group. C) Plasma corticosterone measurement 4 hrs after 250 μg/kg IP LPS treatment. N = 5/group. D) Expression of inflammatory cytokine genes 8 hrs after 250 μg/kg IP LPS treatment. All data are analyzed from ΔCt values and normalized to WT saline group. ** = p<0.01, *** = p<.001 for Bonferroni post-hoc comparisons. # = p<.05, ## = p<.01, ### = p<.001 for interaction effect in Two-way ANOVA. N = 3-4/group. One outlier was removed in the TRIFKO LPS groups due to complete lack of behavioral response to LPS. Results are representative of 2 independent experiments. Figure 1 - figure supplement 1 shows that MyD88 signaling is intact in TRIFKO mice.

CNS inflammation is a hallmark of acute illness responses and cachexia. Therefore, we measured expression of inflammatory cytokine and chemokine genes in the hypothalamus after systemic LPS challenge using qRT-PCR. We found that 8 hrs after 250 μg/kg IP LPS, TRIFKO animals showed attenuated up-regulation of several cytokines and chemokines in the hypothalamus, including *Il6, Ccl2, Ccl5, Cxcl1, Cxcl2,* and *Cxcl10* (Fig. 1d). Alternatively, *Il1β, Tnf, Ifnβ,* and *Cd80* were either similarly upregulated compared to WT LPS-treated mice or not upregulated in either group of LPS-treated mice. It is important to note that basal expression of all cytokines/chemokines was detectable in hypothalami of saline-treated animals.

In order to rule out altered MyD88 signaling as a result of TRIF deletion, we challenged TRIFKO mice with 10 ng ICV IL-1 β. MyD88 is essential for IL-1R signaling, but TRIF is not involved (Muzio, Ni, Feng, & Dixit, 1997). We found that WT and TRIFKO mice had similar anorexia response to ICV IL-1 β (Fig. 1 - figure supplement 1a). While WT IL-1β-treated mice lost more weight than WT saline-treated mice, it was not significantly more than TRIFKO IL-1β-treated mice (Fig. 1 - figure supplement 1b). Lastly, *Myd88* was equally expressed in WT and TRIFKO mice at baseline, and similarly upregulated after IP LPS exposure (Fig. 1 - figure supplement 1b).

### TRIF is important in acute illness response after ICV LPS challenge

To determine the role of TRIF signaling in the CNS after TLR4 activation, we injected LPS directly into the brain lateral ventricles of WT and TRIFKO mice at a dose that has no behavioral effects when injected peripherally (50 ng). While ICV injection of 50 ng LPS caused a significant decrease in cumulative food intake over 62 hrs after injection in both WT and TRIFKO mice, LPS-treated TRIFKO mice consumed more than LPS-treated WT mice starting 36 hrs after treatment (Fig. 2a). Furthermore, TRIFKO mice treated with ICV LPS showed significantly attenuated weight loss compared to WT mice treated with ICV LPS at 24 and 36 hrs after injection (Fig. 2b).

**Figure 2:**
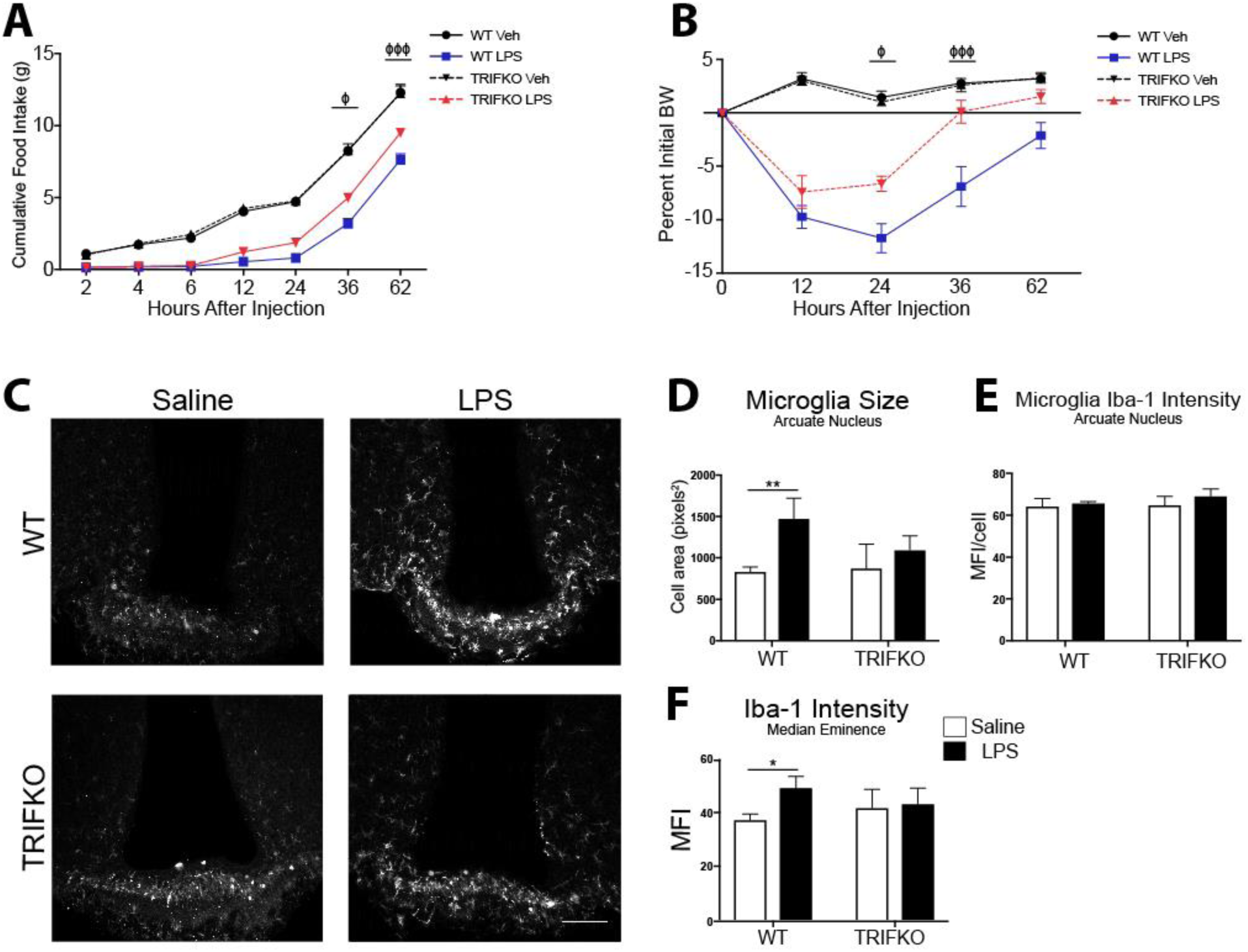
TRIFKO mice have attenuated acute sickness behavior in response to central nervous system LPS exposure. A) Cumulative food intake after 50 ng ICV LPS treatment. Veh = vehicle treatment. B) Body weight change after 50 ng ICV LPS treatment. BW = body weight. Φ = p<0.05, ΦΦΦ = p<0.001 for WT LPS vs. TRIFKO LPS in Bonferroni post-hoc comparisons. N = 5-6/group (See Figure 2 - source data 1 for food intake and body weight data). C) Representative images of Iba-1 immunoreactivity in 200X magnification images of the MBH in WT and TRIFKO mice after either 50 ng ICV LPS or saline. Scale bar = 100 μm. D) Quantification of arcuate nucleus microglia size (area) in pixels^2^ after either ICV LPS or saline. E) Quantification of Iba-1 intensity per microglia in the arcuate nucleus in WT and TRIFKO mice after either ICV LPS or LPS saline. F) Quantification of Iba-1 intensity in the median eminence in WT and TRIFKO mice after either ICV LPS or LPS saline. * = p<0.05, ** = p<0.01. n = 4/group. Data shown are representative or pooled data from 2-5 independent experiments.

### TRIF is required for microglial activation after ICV LPS treatment

TRIF is important for microglia function during states of disease (Hosmane et al., 2012; Sen Lin et al., 2012). However, no studies have investigated the role of TRIF in microglial activation after TLR4 stimulation. Therefore, we quantified microglial activation in the MBH 12 hrs after ICV LPS administration (50 ng) by measuring Iba-1 intensity per cell and cell area. While arcuate nucleus microglia in LPS-treated WT mice showed a significant increase in size compared to saline-treated WT mice, arcuate nucleus microglia in LPS-treated TRIFKO mice did not increase in size compared to saline-treated TRIFKO mice (Fig. 2d). In the arcuate nucleus, Iba-1 intensity per microglia did not increase in the LPS-treated group for either genotype (Fig. 2e). However, overall Iba-1 intensity increased in the median eminence in the WT LPS-treated group compared to the WT saline-treated group, but not in the TRIFKO LPS-treated group compared to the TRIFKO saline-treated group (Fig. 2f).

### TRIF is required for neutrophil recruitment to the brain

Since chemokines comprised the majority of inflammatory transcripts that were less upregulated in TRIFKO mice after LPS exposure, we hypothesized that TRIF is important in immune cell recruitment to the brain. We performed flow cytometry on the brains of WT and TRIFKO mice 12 hrs after 500 ng ICV LPS exposure. We focused on neutrophils because of previous literature showing they are the predominant cell type in the brain after LPS exposure (He et al., 2016). We found that compared to saline-treated WT mice, LPS-treated WT mice had a significantly higher percentage of CD45+ cells in the brain that were neutrophils (Fig. 3a-c). Alternatively, compared to saline-treated TRIFKO mice, LPS-treated TRIFKO mice did not have an increased percentage of CD45+ cells in the brain that were neutrophils (Fig. 3c). There was no increase in T-cells, Ly6C^hi^ monocytes, or Ly6C^low^ monocytes after LPS exposure in either genotype (Fig. 3d).

**Figure 3:**
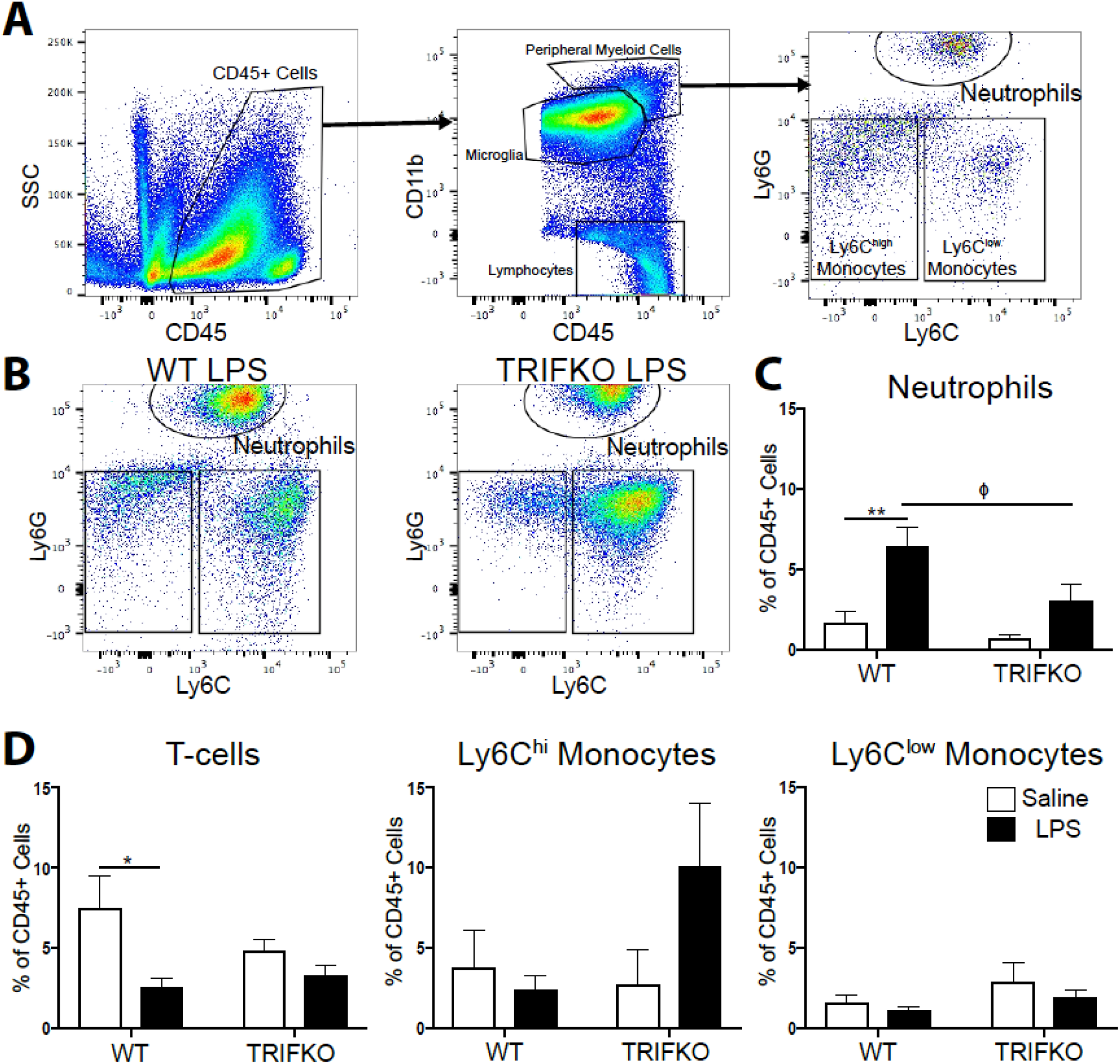
TRIF is required for neutrophil recruitment to the brain after ICV LPS. A) Flow cytometry gating strategy for various immune cell types in the brain from representative WT brain treated with ICV saline. B) Representative flow cytometry plots from WT and TRIFKO brains treated with 500 ng ICV LPS, gated for CD45^high^CD11b+ myeloid cells. C) Quantification of neutrophils in the brain as percentage of total CD45+ in WT and TRIFKO brains treated with either ICV LPS or saline. D) Quantification of CD3+ T-cells, Ly6C^hi^ monocytes, and Ly6C^low^ monocytes in the brain as percentage of total CD45+ in WT and TRIFKO brains treated with either ICV LPS or saline. * = p<0.05, ** = p<0.01. N = 4/group. Φ = p <0.05 for WT LPS vs. TRIFKO LPS in Bonferroni post-hoc comparisons. Data are combined from 4 independent experiments (n = 4). Figure 3 - figure supplement 1 show gating strategy for live singlet cells.

### Mice lacking TRIF have attenuated cancer cachexia

Inflammation is a key component of cachexia (Burfeind et al., 2015), and studies have shown increased production of inflammatory cytokines in the hypothalamus during cancer cachexia (Braun et al., 2011; Michaelis et al., 2017). However, mechanisms of inflammatory signaling in the CNS during cancer cachexia remain unclear. TRIFKO mice inoculated orthotopically with KPC PDAC cells experienced attenuated anorexia compared to WT mice with PDAC (Fig. 4a and b). Furthermore, TRIFKO tumor mice showed attenuated fatigue compared to WT tumor mice (Fig. 4c). WT tumor-bearing mice showed significantly decreased gastrocnemius mass compared to WT sham-operated mice while TRIFKO tumor-bearing mice did not show decreased gastrocnemius mass compared to TRIFKO sham-operated mice (Fig. 4d). These effects on muscle catabolism were further evidenced by the fact that the E3 ubiquitin-ligase system genes *Mafbx* and *Murf1* were upregulated in WT tumor animals compared to WT sham animals, but not significantly upregulated in TRIFKO tumor-bearing animals (Fig. 4e). The same was true for *Foxo1,* a key transcription factor for muscle catabolism (Sandri et al., 2004). In addition, although *Ccl2* was significantly upregulated in the hypothalamus of WT tumor animals, it was not in TRIFKO tumor-bearing animals. Alternatively, compared to WT tumor-bearing animals, *Il1β* was equally upregulated in the hypothalami of TRIFKO tumor-bearing animals, and *Tnf, Il6, Cd80, Cxcl1, Cxcl2,* and *Cxcl10* were not upregulated in WT or TRIFKO tumor-bearing animals. Lastly, although *Ccl5* was less upregulated in TRIFKO tumor-bearing animals compared to WT tumor-bearing animals, this relationship was not significant (Fig. 4f). *Ifnβ* was excluded from analysis due to undetectable expression in several samples.

**Figure 4:**
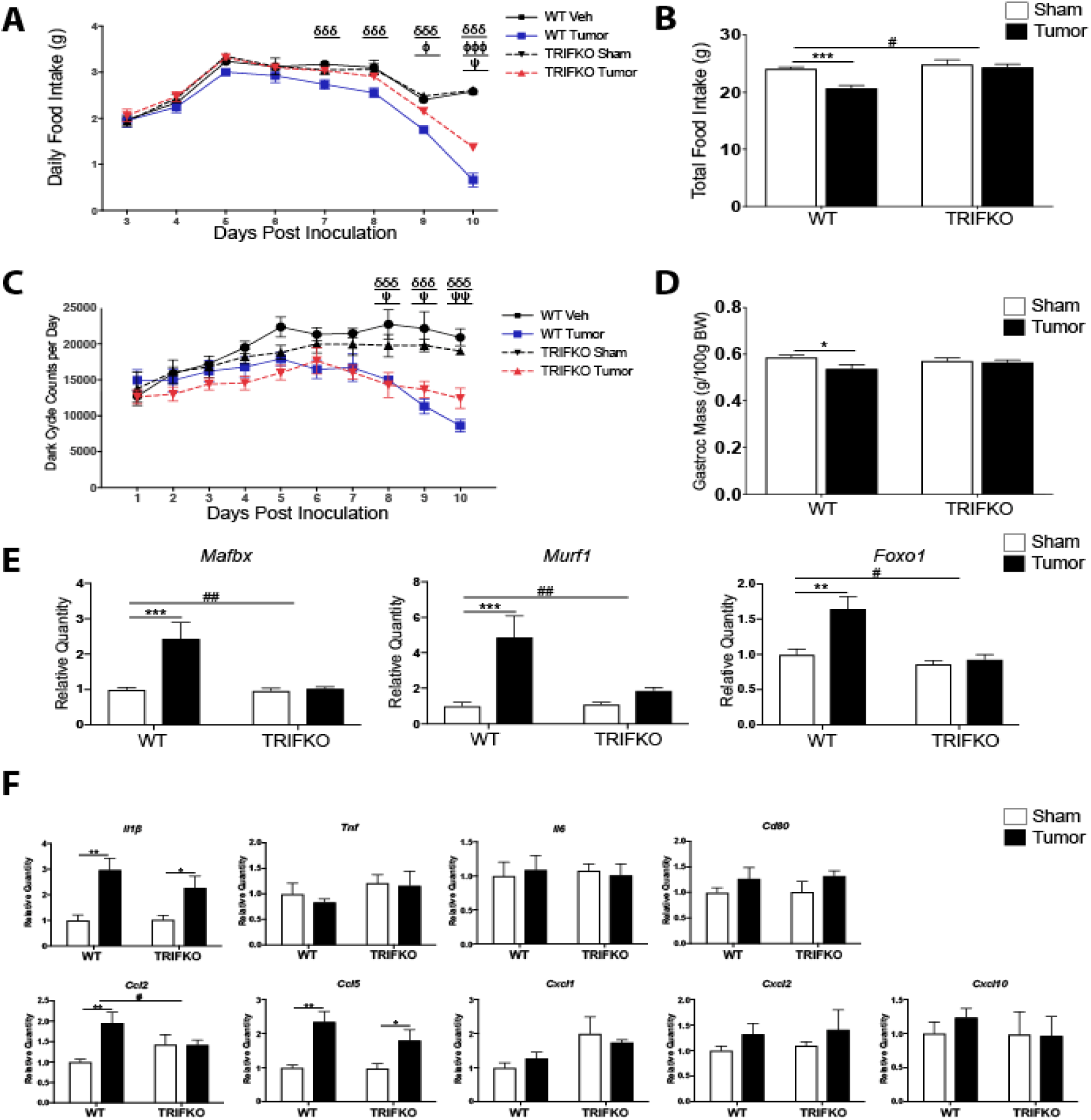
TRIFKO mice have attenuated cancer cachexia. A) Daily food intake after a single orthotopic inoculation of 3e^6^ KPC tumor cells. δδδ = p<0.001 for WT sham vs. WT tumor in Bonferroni post-hoc comparisons. Φ = p<0.05, ΦΦΦ = p<0.001 for WT tumor vs. TRIFKO tumor in Bonferroni post-hoc comparisons. Ψ = p<0.05 for TRIFKO sham vs. TRIFKO tumor in Bonferroni post-hoc comparisons. B) Cumulative food intake 10 days post inoculation. *** = p<0.001 for WT sham vs. WT tumor in Bonferroni post-hoc comparisons. # = p<0.05 for interaction in Two-way ANOVA. C) Movement quantification after inoculation with KPC tumor cells. Movement quantified using a Minimitter system with e-mitter implanted subcutaneously in between shoulder blades. δδδ = p<0.001 for WT sham vs. WT tumor in Bonferroni post-hoc comparisons. Ψ = p<0.05, ΨΨ = p<0.01 for TRIFKO sham vs. TRIFKO tumor in Bonferroni post-hoc comparisons (See Figure 4 - source data 1 for raw minimitter data). D) Muscle catabolism determined by gastrocnemius mass. Mass of dissected left and right gastrocnemius was averaged and then divided by initial body weight for normalization (See Figure 4 - source data 2 for food intake and body weight data). * = p<0.05 for WT sham vs. WT tumor in Bonferroni post-hoc comparisons. E) qRT-PCR analysis of muscle catabolism genes in gastrocnemius. Expression level for all groups was normalized to WT sham. ** = p<0.01, *** = p<0.001 for WT sham vs. WT tumor in Bonferroni post-hoc comparisons. # = p<0.05, ## = p<0.01 for interaction effect in Two-way ANOVA. F) Expression of inflammatory cytokine genes 10 days after orthotopic inoculation with KPC tumor cells. All data are analyzed from ΔCt values and normalized to WT sham group. * = p<0.05, ** = p<0.01, *** = p<0.001 for WT sham vs. WT tumor in Bonferroni post-hoc comparisons. # = p<0.05, ## = p<0.01 for interaction effect in Two-way ANOVA. N = 5/group for all nexperiments. Data are representative from 3 independent experiments.

## Discussion

We investigated the role of TRIF in acute sickness behavior after LPS exposure and in a model of pancreatic cancer cachexia. These studies demonstrated that TRIFKO mice experienced attenuated sickness behavior after peripheral or central LPS exposure. Furthermore, TRIFKO mice experienced attenuated cachexia during PDAC, including decreased anorexia, fatigue, muscle catabolism, and hypothalamic inflammation relative to WT counterparts. These results indicate that TRIF is an important mediator of inflammation-driven sickness behavior, and should be considered during the development of anti-inflammatory therapies for cachexia.

Several studies investigated the role of MyD88 in sickness behavior (Braun et al., 2012; Yamawaki, Kimura, Hosoi, & Ozawa, 2010; Zhu, Levasseur, Michaelis, Burfeind, & Marks, 2016), yet evidence suggests that MyD88-independent signaling pathways are important in CNS immune activation (Hosmane et al., 2012; Menasria et al., 2013). While the role of TRIF signaling in acute sickness behavior during viral infection was investigated previously (Gibney, McGuinness, Prendergast, Harkin, & Connor, 2013; Murray et al., 2015; Zhu et al., 2016), this is the first to investigate the role of TRIF in acute sickness behavior after LPS exposure and during cachexia. After systemic challenge with LPS, TRIFKO mice experienced attenuated anorexia and weight loss compared to WT mice. This coincided with an attenuated increase in serum corticosterone, implicating TRIF as a key player in stress response.

In addition to attenuated acute sickness behavior after systemic LPS exposure, TRIFKO mice experienced attenuated anorexia and weight loss after ICV LPS administration. Interestingly, TRIFKO mice showed similar weight loss 12 hours after LPS administration, yet recovered more rapidly than WT mice, reaching baseline body weight 36 hours after injection. These results suggest that in the CNS, MyD88 may drive initial sickness response after TLR4 activation, whereas TRIF signaling may be involved in maintaining inflammation and subsequent sickness response.

In the brain, microglia express TRIF at basal levels, and this expression is enhanced by various CNS insults (S. Lin et al., 2012; Y. Wang et al., 2013). Review of the published online database (http://web.stanford.edu/group/barreslab/brainrnaseq.html) confirms that the basal expression of TRIF (*Ticam1*) is predominantly found in microglia (Zhang et al., 2014). We found that TRIFKO mice had attenuated microglial activation 12 hrs after ICV LPS administration. This is in agreement with previous studies showing that that TRIF expression in microglia is required for normal inflammatory activation and phagocytosis in response to neuronal injury (Hosmane et al., 2012; Sen Lin et al., 2012). Furthermore, in a murine model of intracerebral hemorrhage, TRIFKO mice showed attenuated neurologic disability and neuroinflammation (Sen Lin et al., 2012). In addition, TLR2 activation in hypothalamic microglia was shown to generate sickness responses (Jin et al., 2016a), and TRIF is now known to be linked to TLR2 signaling (Nilsen et al., 2015; Petnicki-Ocwieja et al., 2013). Therefore, our data, in addition to previous findings, suggest TRIF is important in microglial activation during states of inflammation, which is important in driving subsequent functional and behavioral response.

We found that TRIFKO mice experienced attenuated hypothalamic inflammation after systemic LPS exposure. Interestingly, amongst the differentially regulated transcripts between WT and TRIFKO animals, there was a predominance of chemokine mRNAs (*Ccl2, Ccl5, Cxcl1*, *Cxcl10*). Previous studies showed that LPS exposure results in peripheral immune cell recruitment to the brain (He et al., 2016) and that infiltrating immune cells in the brain drive sickness behavior (D’Mello, Le, & Swain, 2009). Based on these data, we investigated whether TRIFKO mice had decreased immune cell infiltration into the brain after ICV LPS exposure and found that TRIF was required for neutrophil recruitment. While TRIF is known to be important for neutrophil recruitment to the lungs (Liu et al., 2016), this is the first study to implicate TRIF in neutrophil recruitment to the brain. This presents a novel mechanism that can be applied to several pathologies, including CNS infection, cancer, and stroke. Furthermore, no studies have investigated whether neutrophils are important in sickness behavior or cachexia.

When inflammation is maintained, acute sickness behavior transforms into cachexia, a maladaptive condition associated with increased mortality and decreased quality of life during numerous chronic diseases (Bachmann et al., 2008; Lainscak et al., 2007; Wesseltoft-Rao et al., 2015). While inflammation is critical for cachexia, mechanisms of inflammatory signaling important for this syndrome remain unclear. We found that in a mouse model of PDAC-associated cachexia, TRIFKO mice experienced attenuated anorexia, fatigue, muscle catabolism, and hypothalamic inflammation compared to WT mice. It is important to note that differences between WT and TRIFKO mice only emerged 9-10 days after tumor inoculation, suggesting that TRIF is important in later stages of cachexia. Therefore, it is possible that the MyD88 pathway dominates in the initiation stage of cachexia. Ruud et al. reported that mice lacking MyD88 in immune cells experienced attenuated anorexia and muscle catabolism during cancer cachexia (Ruud et al., 2013). However, their results were similar to ours in that MyD88^ΔMX1Cre^ tumor-bearing animals experienced attenuated anorexia compared to WT tumor-bearing animals at 10 days post inoculation, which was at the end-stage of cachexia. *MX1* is expressed mainly in cells of the myeloid lineage, suggesting MyD88 in other cell types, or different inflammatory pathways may be responsible for initiation of cachexia.

The main limitation of the present study is the lack of cell specificity in global TRIFKO experiments. Future studies are needed to identify the critical cell type involved in TRIF-mediated sickness behavior and cachexia. Another limitation is the fact that we performed analysis of cachexia using only one mouse model of cancer cachexia. Caution is warranted when applying our results to other types of cachexia (heart failure, cirrhosis, untreated HIV, other types of cancer, etc.). However, this model is extensively characterized (Michaelis et al., 2017), and recapitulates all of the cardinal features of cachexia seen in humans. Furthermore, it avoids many of the shortcomings in other mouse models of cachexia, including: multiple clones with variable cachexia (Kir et al., 2014), cachexia driven by only a single cytokine (Talbert, Metzger, He, & Guttridge, 2014), and requiring advanced surgical techniques to induce cachexia (DeBoer, 2009).

In conclusion, we report that TRIF is important in acute sickness behavior and cachexia. These results show that TRIF-dependent mechanisms should be considered when developing therapeutic targets for cachexia. Future studies are needed to identify the important cell types involved in TRIF signaling during acute illness response and cachexia.

## Materials and Methods

### Animals

Male and female 20–25-g WT C57BL/6J (stock no. 000664) and TRIFKO (Trif^Lps2^, stock no. 005037) mice were obtained from The Jackson Laboratory. Mice were between 7 and 12 weeks of age at time of experiment. All animals were maintained at 27°C on a normal 12:12 hr light/dark cycle and provided *ad libitum* access to water and food. Experiments were conducted in accordance with the National Institutes of Health Guide for the Care and Use of Laboratory Animals, and approved by the Animal Care and Use Committee of Oregon Health and Science University.

### Intracerebroventricular Cannulation and Injections

Mice were anesthetized under isoflurane and placed on a stereotactic alignment instrument (Kopf Instruments, CA). 26-gauge lateral ventricle cannulas were placed at -1.0 mm X, -0.5 mm Y, and -2.25 mm Z relative to bregma. Injections were given in 2 μl total volume. LPS (from *Escherichia coli*, O555:B5, Sigma Aldrich, St. Louis, MO) was dissolved in normal saline with 0.5% bovine serum albumin.

### Nocturnal Feeding Studies

Animals were transferred to clean cages and injected with ICV (50 ng) or IP (250 μg) LPS 1 h prior to lights off. At 2, 6, 12, 24, 36 and 48 hrs after the onset of the dark cycle, food was weighed and returned to the cage. Body weight was recorded at 12, 24 and 48 hrs.

### Plasma Corticosterone Measurement

Plasma corticosterone levels were measured by RIA (MP Biomedicals, Valiant, Yantai, China) according to the manufacturer’s instructions. Animals were anesthetized with a lethal dose of a ketamine/xylazine/acetapromide 4 hrs after IP LPS administration. Blood was obtained by cardiac puncture, anticoagulated with EDTA and separated by centrifugation. Plasma was stored at −80°C until analysis.

### Quantitative Real-Time PCR

Prior to tissue extraction, mice were euthanized with a lethal dose of a ketamine/xylazine/acetapromide and sacrificed. Hypothalamic blocks were dissected, snap frozen, and stored in −80 °C until analysis. Hypothalamic RNA was extracted using an RNeasy mini kit (Qiagen, Hilden, Germany) according to the manufacturer’s instructions. cDNA was transcribed using TaqMan reverse transcription reagents and random hexamers according to the manufacturer’s instructions. PCR reactions were run on an ABI 7300 (Applied Biosystems), using TaqMan universal PCR master mix with the following TaqMan mouse gene expression assays: *18s* (Mm04277571_s1), *Il1b* (Mm00434228_m1), *Tnf* (Mm00443258_m1), *Il6* (Mm01210732_g1), *Cd80* (Mm00711660_m1), *Myd88* (Mm00440338_m1), *Ifnb1* (Mm00439552_s1), *Ccl2* (Mm99999056_m1), *Ccl5* (Mm01302427_m1), *Cxcl1* (Mm04207460_m1), *Cxcl2* (Mm00436450_m1), *Cxcl10* (Mm00445235_m1), *Gapdh* (Mm99999915_g1), *Mafbx* (Mm00499518_m1), *Murf1* (Mm01185221_m1), and *Foxo1* (Mm00490672_m1).

Relative expression was calculated using the ΔΔCt method and normalized to vehicle treated or sham control. Statistical analysis was performed on the normally distributed ΔCt values.

### Immunohistochemistry

Mice were anesthetized using a ketamine/xylazine/acetapromide cocktail and sacrificed by transcardial perfusion fixation with 15 mL ice cold 0.01 M PBS followed by 25 mL 4% paraformaldehyde (PFA) in 0.01 M PBS. Brains were post-fixed in 4% PFA overnight at 4°C and cryoprotected in 20% sucrose for 24 hrs at 4°C before being stored at −80°C until used for immunohistochemistry. Immunofluorescence histochemistry was performed as described below. Free-floating sections were cut at 30 μm from perfused brains using a sliding microtome (Leica SM2000R, Leica Microsystems, Wetzlar, Germany). Hypothalamic sections were collected from the division of the optic chiasm (bregma −1.0 mm) caudally through the mammillary bodies (bregma −3.0 mm). The sections were incubated for 30 min at room temperature in blocking reagent (5% normal donkey serum in 0.01 M PBS and 0.3% Triton X-100). After the initial blocking step, the sections were incubated in rabbit anti-mouse Iba-1 (1:500, DAKO) in blocking reagent for 24 hrs at 4°C, followed by incubation in donkey anti-rabbit Alexa 555 (1:1000) for 2 hrs at room temperature. Between each stage, the sections were washed thoroughly with 0.01 M PBS. Sections were mounted onto gelatin-coated slides and coverslipped using Prolong Gold Antifade media with DAPI (Thermofisher, Waltham, MA).

### Microglia Activation Quantification

Microglia activation in the MBH was quantified using Fiji (ImageJ, NIH, Bethesda, MD). The MBH was defined as the region surrounding the third ventricle at the base of the brain, starting rostrally at the end of the optic chiasm when the arcuate nucleus appears (−1.22 mm from bregma) and ending caudally at the mammillary body (−2.70 mm from bregma). Images were acquired using the 20X objective (na=0.8, step size=1 μm), with the base of the MBH positioned at the very bottom of the field of view (FOV) and the third ventricle at the center of the FOV. Care was taken to exclude the meninges so as to avoid analysis of meningeal macrophages. Images were 2048 × 2048 pixels, with a pixel size of 0.315 μm. Images were acquired as 8-bit RGB TIFF images. 3-10 MBH images per animal were acquired and analyzed.

After image acquisition, TIFF images were uploaded to Fiji and converted to 8-bit greyscale images. After thresholding, microglia were identified using the Analyze Particle function, which measured mean Iba-1 fluorescent intensity per cell and cell area. Iba-1 fluorescent intensity and cell size was measured for each microglia in the arcuate nucleus. Due to the density of microglia in the median eminence (ME), the software was unable to differentiate individual cells. As such, overall Iba-1 fluorescent intensity was measured to quantify microglia activation in the ME.

### Flow Cytometry

12 hrs after 500 ng ICV LPS administration, mice were anesthetized using a ketamine/xylazine/acetapromide cocktail and perfused with 15 mL ice cold 0.01 M PBS to remove circulating leukocytes. After perfusion, brains were extracted and minced in a digestion solution containing 1 mg/mL type II collagenase (Sigma) and 1% DNAse (Sigma) in RPMI, then placed in a 37°C incubator for 1 hr. After digestion, myelin was removed via using 30% percoll in RPMI. Isolated cells were washed with RPMI, incubated in Fc block for 5 min, then stained with the following antibodies (all rat anti-mouse from BioLegend, except for Live/Dead) (BioLegend, San Diego, CA): anti-CD45 PerCP/Cy5.5 (1:400), anti-CD11b APC (1:800), anti-Ly6C PerCP (1:100), anti-Ly6G PE/Cy7 (1:800), anti-CD3 PE (1:100), and Live/Dead fixable aqua (1:200, Thermofisher). Flow cytometry was conducted using a Fortessa analytic flow cytometer (BD Biosciences, NJ), and analysis was performed on FlowJo V10 software (FlowJo, Ashland, OR). Cells were gated on LD, SSC singlet, and FSC singlet (Fig. 3 - figure supplement 1). Leukocytes were then defined as CD45+ cells and identified as either peripheral myeloid cells (CD45^high^CD11b+) or lymphocytes (CD45^high^CD11b−). From peripheral myeloid cells Ly6C^low^ monocytes (Ly6C^low^Ly6G−), Ly6C^high^ monocytes (Ly6C^high^Ly6G−), and neutrophils (Ly6C^mid^Ly6G+) were identified. From lymphocytes, CD3+ cells were identified as T-cells.

### KPC Cancer Cachexia Model

Our lab generated a mouse model of pancreatic ductal adenocarcinoma (PDAC) – associated cachexia by injection of murine-derived KPC PDAC cells (originally provided by Dr. Elizabeth Jaffee from Johns Hopkins) (Michaelis et al., 2017). These cells are derived from tumors in mice with <u>K</u>RAS^G12D^ and <u>T</u>P53^R172H^ deleted via the PDX-1-<u>C</u>re driver (Foley et al., 2015). Cells were maintained in RPMI supplemented with 10% heat-inactivated FBS, and 50 U/mL penicillin/streptomycin (Gibco, Thermofisher), in incubators maintained at 37°C and 5% CO2. In the week prior to tumor implantation, animals were transitioned to individual housing to acclimate to experimental conditions. Animal food intake and body weight were monitored daily. Mice were inoculated orthotopically with 3 million KPC tumor cells in 40 μL PBS into the tail of the pancreas (Chai, Kim-Fuchs, Angst, & Sloan, 2013). Sham-operated animals received heat-killed cells in the same volume. NMR measurements were taken at the beginning of the study for covariate adaptive randomization of tumor and sham groups to ensure equally distributed weight and body composition. Body temperature and voluntary home cage locomotor activity were measured via MiniMitter tracking devices (Starr Life Sciences, Oakmont, PA). Mice were implanted 7 days prior to tumor implantation with MiniMitter transponders in the intrascapular subcutaneous space. Using these devices, movement counts in *x*-axis, *y*-axis, and *z*-axis were recorded in 5 min intervals.

### Statistical Analysis

Data are expressed as means ± SEM. Statistical analysis was performed with Prism 7.0 software (Graphpad Software Corp, La Jolla, CA). All data were analyzed with a Two-way ANOVA followed with *post hoc* analysis with a Bonferroni *post hoc* test or Student’s *t* test as appropriate. For all analyses, significance was assigned at the level of p< 0.05.

### Competing Interests

The authors have no competing interests to report.

## Supplemental Table and Figure Legends

**Figure 1 - Figure Supplement 1:**
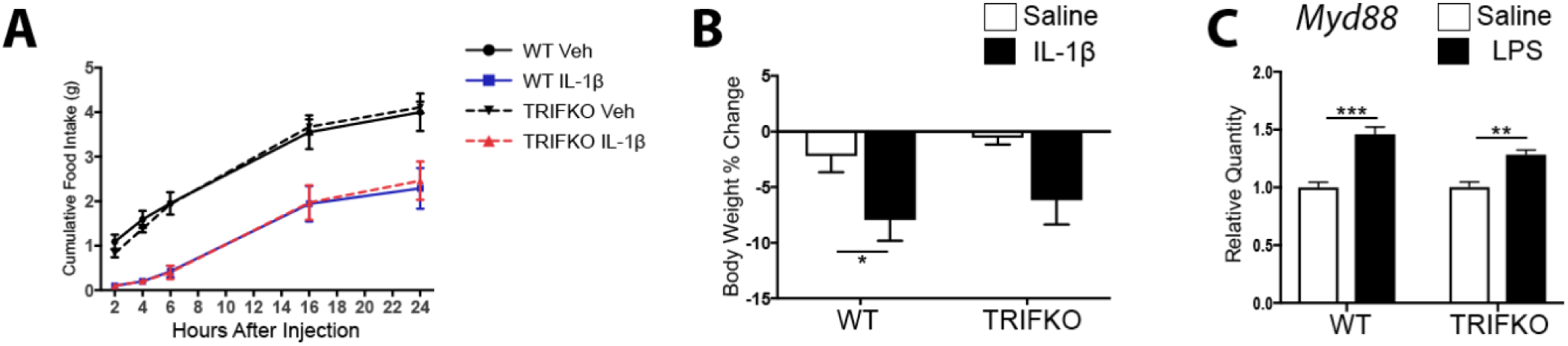
MyD88 signaling is intact in TRIFKO mice. A)Cumulative food intake after 10 ng ICV IL-1β in artificial CSF or vehicle (artificial CSF only) treatment. B) Body weight change from initial after 10 ng ICV IL-1 β in artificial CSF or vehicle (artificial CSF only) treatment. N=6-8/group. C) qRT-PCR for *Myd88* gene expression in WT and TRIFKO mice. LPS = 8 hrs after 250 μg/kg IP LPS treatment. N = 3-4/group. * = p<0.05, ** = p<0.01, *** = p<0.001 for two-way ANOVA Bonferroni post-hoc testing.

**Figure 3 - Figure Supplement 1:**
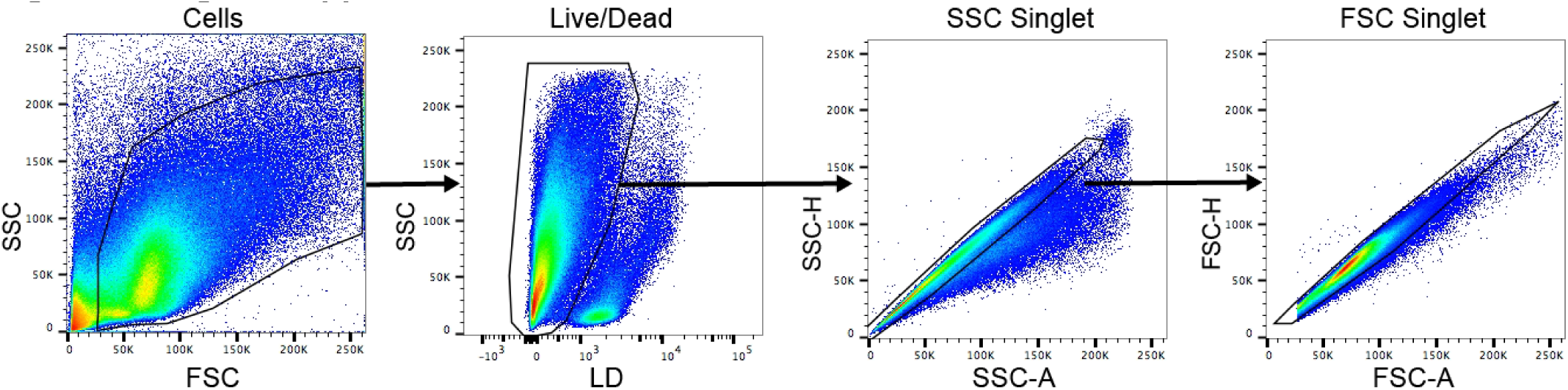
Flow cytometry gating strategy. Representative plots from FlowJo V10 software. SSC = side scatter. FSC = forward scatter. A = area. H = height. LD = Live/Dead.

## References Cited

Anker, S. D., Ponikowski, P., Varney, S., Chua, T. P., Clark, A. L., Webb-Peploe, K. M., Harrington, D., Kox, W. J., Poole-Wilson, P. A., & Coats, A. J. (1997). Wasting as independent risk factor for mortality in chronic heart failure. Lancet, 349(9058), 1050–1053.

Argiles, J. M., Anker, S. D., Evans, W. J., Morley, J. E., Fearon, K. C., Strasser, F., Muscaritoli, M., & Baracos, V. E. (2010). Consensus on cachexia definitions. J Am Med Dir Assoc, 11(4), 229–230. doi:10.1016/j.jamda.2010.02.004

Bachmann, J., Heiligensetzer, M., Krakowski-Roosen, H., Buchler, M. W., Friess, H., & Martignoni, M. E. (2008). Cachexia worsens prognosis in patients with resectable pancreatic cancer. J Gastrointest Surg, 12(7), 1193–1201. doi:10.1007/s11605-008-0505-z

Bluthé, R.-M., Michaud, B., Poli, V., & Dantzer, R. (2000). Role of IL-6 in cytokine-induced sickness behavior: a study with IL-6 deficient mice. Physiol Behav, 70(3-4), 367–373. doi:https://doi.org/10.1016/S0031-9384(00)00269-9

Bodnar, R. J., Pasternak, G. W., Mann, P. E., Paul, D., Warren, R., & Donner, D. B. (1989). Mediation of anorexia by human recombinant tumor necrosis factor through a peripheral action in the rat. Cancer Res, 49(22), 6280–6284.

Braun, T. P., Grossberg, A. J., Veleva-Rotse, B. O., Maxson, J. E., Szumowski, M., Barnes, A. P., & Marks, D. L. (2012). Expression of myeloid differentiation factor 88 in neurons is not requisite for the induction of sickness behavior by interleukin-1beta. J Neuroinflammation, 9, 229. doi:10.1186/1742-2094-9-229

Braun, T. P., Zhu, X., Szumowski, M., Scott, G. D., Grossberg, A. J., Levasseur, P. R., Graham, K., Khan, S., Damaraju, S., Colmers, W. F., Baracos, V. E., & Marks, D. L. (2011). Central nervous system inflammation induces muscle atrophy via activation of the hypothalamic-pituitary-adrenal axis. J Exp Med, 208(12), 2449–2463. doi:10.1084/jem.20111020

Burfeind, K. G., Michaelis, K. A., & Marks, D. L. (2015). The central role of hypothalamic inflammation in the acute illness response and cachexia. Semin Cell Dev Biol. doi:10.1016/j.semcdb.2015.10.038

Chai, M. G., Kim-Fuchs, C., Angst, E., & Sloan, E. K. (2013). Bioluminescent orthotopic model of pancreatic cancer progression. J Vis Exp(76). doi:10.3791/50395

D’Mello, C., Le, T., & Swain, M. G. (2009). Cerebral microglia recruit monocytes into the brain in response to tumor necrosis factoralpha signaling during peripheral organ inflammation. J Neurosci, 29(7), 2089–2102. doi:10.1523/jneurosci.3567-08.2009

Dantzer, R., BluthÉ, R.-M., LayÉ, S., Bret-Dibat, J.-L., Parnet, P., & Kelley, K. W. (1998). Cytokines and Sickness Behavior. Ann N Y Acad Sci, 840(1), 586–590. doi:10.1111/j.1749-6632.1998.tb09597.x

DeBoer, M. D. (2009). Animal models of anorexia and cachexia. Expert opinion on drug discovery, 4(11), 1145–1155. doi:10.1517/17460440903300842

Elmquist, J. K., Scammell, T. E., Jacobsen, C. D., & Saper, C. B. (1996). Distribution of Fos-like immunoreactivity in the rat brain following intravenous lipopolysaccharide administration. J Comp Neurol, 371(1), 85–103.

Evans, W. J., Morley, J. E., Argiles, J., Bales, C., Baracos, V., Guttridge, D., Jatoi, A., Kalantar-Zadeh, K., Lochs, H., Mantovani, G., Marks, D., Mitch, W. E., Muscaritoli, M., Najand, A., Ponikowski, P., Rossi Fanelli, F., Schambelan, M., Schols, A., Schuster, M., Thomas, D., Wolfe, R., & Anker, S. D. (2008). Cachexia: a new definition. Clin Nutr, 27(6), 793–799. doi:10.1016/j.clnu.2008.06.013

Fearon, K., Strasser, F., Anker, S. D., Bosaeus, I., Bruera, E., Fainsinger, R. L., Jatoi, A., Loprinzi, C., MacDonald, N., Mantovani, G., Davis, M., Muscaritoli, M., Ottery, F., Radbruch, L., Ravasco, P., Walsh, D., Wilcock, A., Kaasa, S., & Baracos, V. E. (2011). Definition and classification of cancer cachexia: an international consensus. Lancet Oncol, 12(5), 489–495. doi:10.1016/S1470-2045(10)70218-7

Feng, Y., Zou, L., Zhang, M., Li, Y., Chen, C., & Chao, W. (2011). MyD88 and Trif signaling play distinct roles in cardiac dysfunction and mortality during endotoxin shock and polymicrobial sepsis. Anesthesiology, 115(3), 555–567. doi:10.1097/ALN.0b013e31822a22f7

Foley, K., Rucki, A. A., Xiao, Q., Zhou, D., Leubner, A., Mo, G., Kleponis, J., Wu, A. A., Sharma, R., Jiang, Q., Anders, R. A., Iacobuzio-Donahue, C. A., Hajjar, K. A., Maitra, A., Jaffee, E. M., & Zheng, L. (2015). Semaphorin 3D autocrine signaling mediates the metastatic role of annexin A2 in pancreatic cancer. SciSignal, 8(388), ra77. doi:10.1126/scisignal.aaa5823

Gibney, S. M., McGuinness, B., Prendergast, C., Harkin, A., & Connor, T. J. (2013). Poly I:C-induced activation of the immune response is accompanied by depression and anxiety-like behaviours, kynurenine pathway activation and reduced BDNF expression. Brain Behav Immun, 28(Supplement C), 170–181. doi:https://doi.org/10.1016/j.bbi.2012.11.010

Gong, S., Miao, Y.-L., Jiao, G.-Z., Sun, M.-J., Li, H., Lin, J., Luo, M.-J., & Tan, J.-H. (2015). Dynamics and Correlation of Serum Cortisol and Corticosterone under Different Physiological or Stressful Conditions in Mice. PLoS One, 10(2), e0117503. doi:10.1371/journal.pone.0117503

Grossberg, A. J., Zhu, X., Leinninger, G. M., Levasseur, P. R., Braun, T. P., Myers, M. G., Jr., & Marks, D. L. (2011). Inflammation-induced lethargy is mediated by suppression of orexin neuron activity. J Neurosci, 31(31), 11376–11386. doi:10.1523/JNEUROSCI.2311-11.2011

Hanke, M. L., & Kielian, T. (2011). Toll-like receptors in health and disease in the brain: mechanisms and therapeutic potential. Clinical science (London, England: 1979), 121(9), 367–387. doi:10.1042/CS20110164

He, H., Geng, T., Chen, P., Wang, M., Hu, J., Kang, L., Song, W., & Tang, H. (2016). NK cells promote neutrophil recruitment in the brain during sepsis-induced neuroinflammation. Scientific Reports, 6, 27711. doi:10.1038/srep27711

Hosmane, S., Tegenge, M. A., Rajbhandari, L., Uapinyoying, P., Kumar, N. G., Thakor, N., & Venkatesan, A. (2012). TRIF mediates microglial phagocytosis of degenerating axons. The Journal of Neuroscience, 32(22), 7745–7757. doi:10.1523/JNEUROSCI.0203-12.2012

Jin, S., Kim, J. G., Park, J. W., Koch, M., Horvath, T. L., & Lee, B. J. (2016a). Hypothalamic TLR2 triggers sickness behavior via a microglia-neuronal axis. 6, 29424. doi:10.1038/srep29424 https://www.nature.com/articles/srep29424-supplementary-information

Jin, S., Kim, J. G., Park, J. W., Koch, M., Horvath, T. L., & Lee, B. J. (2016b). Hypothalamic TLR2 triggers sickness behavior via a microglia-neuronal axis. Scientific Reports, 6, 29424. doi:10.1038/srep29424 https://www.nature.com/articles/srep29424-supplementary-information

Kir, S., White, J. P., Kleiner, S., Kazak, L., Cohen, P., Baracos, V. E., & Spiegelman, B. M. (2014). Tumour-derived PTH-related protein triggers adipose tissue browning and cancer cachexia. Nature, 513(7516), 100–104. doi:10.1038/nature13528

Konsman, J. P., Tridon, V., & Dantzer, R. (2000). Diffusion and action of intracerebroventricularly injected interleukin-1 in the CNS. Neuroscience, 101(4), 957–967.

Kotler, D. P., Tierney, A. R., Wang, J., & Pierson, R. N. J. (1989). Magnitude of body-cell-mass depletion and the timing of death from wasting in AIDS. Am. J. Clin. Nutr., 50(3), 444–447.

Laflamme, N., & Rivest, S. (1999). Effects of systemic immunogenic insults and circulating proinflammatory cytokines on the transcription of the inhibitory factor kappaB alpha within specific cellular populations of the rat brain. J Neurochem, 73(1), 309–321.

Lainscak, M., Podbregar, M., & Anker, S. D. (2007). How does cachexia influence survival in cancer, heart failure and other chronic diseases? Curr Opin Support Palliat Care, 1(4), 299–305.

Lin, S., Yin, Q., Zhong, Q., Lv, F.-L., Zhou, Y., Li, J.-Q., Wang, J.-Z., Su, B.-y., & Yang, Q.-W. (2012). Heme activates TLR4-mediated inflammatory injury via MyD88/TRIF signaling pathway in intracerebral hemorrhage. J Neuroinflammation, 9(1), 46. doi:10.1186/1742-2094-9-46

Lin, S., Yin, Q., Zhong, Q., Lv, F. L., Zhou, Y., Li, J. Q., Wang, J. Z., Su, B. Y., & Yang, Q. W. (2012). Heme activates TLR4-mediated inflammatory injury via MyD88/TRIF signaling pathway in intracerebral hemorrhage. J Neuroinflammation, 9, 46. doi:10.1186/1742-2094-9-46

Liu, Y., Gu, Y., Han, Y., Zhang, Q., Jiang, Z., Zhang, X., Huang, B., Xu, X., Zheng, J., & Cao, X. (2016). Tumor Exosomal RNAs Promote Lung Pre-metastatic Niche Formation by Activating Alveolar Epithelial TLR3 to Recruit Neutrophils. Cancer Cell, 30(2), 243–256. doi:10.1016/j.ccell.2016.06.021

Medzhitov, R., Preston-Hurlburt, P., Kopp, E., Stadlen, A., Chen, C., Ghosh, S., & Janeway, C. A., Jr. (1998). MyD88 is an adaptor protein in the hToll/IL-1 receptor family signaling pathways. Mol Cell, 2(2), 253–258.

Menasria, R., Boivin, N., Lebel, M., Piret, J., Gosselin, J., & Boivin, G. (2013). Both TRIF and IPS-1 adaptor proteins contribute to the cerebral innate immune response against HSV-1 infection. J Virol. doi:10.1128/JVI.00591-13

Michaelis, K. A., Zhu, X., Burfeind, K. G., Krasnow, S. M., Levasseur, P. R., Morgan, T. K., & Marks, D. L. (2017). Establishment and characterization of a novel murine model of pancreatic cancer cachexia. J Cachexia Sarcopenia Muscle, 8(5), 824–838. doi:10.1002/jcsm.12225

Morgan, J. I., & Curran, T. (1986). Role of ion flux in the control of c-fos expression. Nature, 322(6079), 552–555. doi:10.1038/322552a0

Murray, C., Griffin, É. W., O’Loughlin, E., Lyons, A., Sherwin, E., Ahmed, S., Stevenson, N. J., Harkin, A., & Cunningham, C. (2015). Interdependent and independent roles of type I interferons and IL-6 in innate immune, neuroinflammatory and sickness behaviour responses to systemic poly I:C. Brain Behav Immun, 48, 274–286. doi:10.1016/j.bbi.2015.04.009

Muzio, M., Ni, J., Feng, P., & Dixit, V. M. (1997). IRAK (Pelle) family member IRAK-2 and MyD88 as proximal mediators of IL-1 signaling. Science, 278(5343), 1612–1615.

Nilsen, N. J., Vladimer, G. I., Stenvik, J., Orning, M. P., Zeid-Kilani, M. V., Bugge, M., Bergstroem, B., Conlon, J., Husebye, H., Hise, A. G., Fitzgerald, K. A., Espevik, T., & Lien, E. (2015). A role for the adaptor proteins TRAM and TRIF in toll-like receptor 2 signaling. J Biol Chem, 290(6), 3209–3222. doi:10.1074/jbc.M114.593426

Petnicki-Ocwieja, T., Chung, E., Acosta, D. I., Ramos, L. T., Shin, O. S., Ghosh, S., Kobzik, L., Li, X., & Hu, L. T. (2013). TRIF Mediates Toll-Like Receptor 2-Dependent Inflammatory Responses to Borrelia burgdorferi. Infect Immun, 81(2), 402–410. doi:10.1128/IAI.00890-12

Ruud, J., Wilhelms, D. B., Nilsson, A., Eskilsson, A., Tang, Y.-J., Ströhle, P., Caesar, R., Schwaninger, M., Wunderlich, T., Bäckhed, F., Engblom, D., & Blomqvist, A. (2013). Inflammation- and tumor-induced anorexia and weight loss require MyD88 in hematopoietic/myeloid cells but not in brain endothelial or neural cells. The FASEB Journal, 27(5), 1973–1980. doi:10.1096/fj.12-225433

Sandri, M., Sandri, C., Gilbert, A., Skurk, C., Calabria, E., Picard, A., Walsh, K., Schiaffino, S., Lecker, S. H., & Goldberg, A. L. (2004). Foxo transcription factors induce the atrophy-related ubiquitin ligase atrogin-1 and cause skeletal muscle atrophy. Cell, 117(3), 399–412.

Sonti, G., Ilyin, S. E., & Plata-Salaman, C. R. (1996). Anorexia induced by cytokine interactions at pathophysiological concentrations. Am J Physiol, 270(6 Pt 2), R1394–1402.

Talbert, E. E., Metzger, G. A., He, W. A., & Guttridge, D. C. (2014). Modeling human cancer cachexia in colon 26 tumor-bearing adult mice. Journal of Cachexia, Sarcopenia and Muscle, 5(4), 321–328. doi:10.1007/s13539-014-0141-2

Tisdale, M. J. (2002). Cachexia in cancer patients. Nat Rev Cancer, 2(11), 862–871.

Wang, A. Y., Sea, M. M., Tang, N., Sanderson, J. E., Lui, S. F., Li, P. K., & Woo, J. (2004). Resting energy expenditure and subsequent mortality risk in peritoneal dialysis patients. J Am Soc Nephrol, 15(12), 3134–3143. doi:10.1097/01.asn.0000144206.29951.b2

Wang, Y., He, H., Li, D., Zhu, W., Duan, K., Le, Y., Liao, Y., & Ou, Y. (2013). The role of the TLR4 signaling pathway in cognitive deficits following surgery in aged rats. Mol Med Rep, 7(4), 1137–1142. doi:10.3892/mmr.2013.1322

Wesseltoft-Rao, N., Hjermstad, M. J., Ikdahl, T., Dajani, O., Ulven, S. M., Iversen, P. O., & Bye, A. (2015). Comparing two classifications of cancer cachexia and their association with survival in patients with unresected pancreatic cancer. Nutr Cancer, 67(3), 472–480. doi:10.1080/01635581.2015.1004728

Wisse, B. E., Ogimoto, K., Tang, J., Harris, M. K., Jr., Raines, E. W., & Schwartz, M. W. (2007). Evidence that lipopolysaccharide-induced anorexia depends upon central, rather than peripheral, inflammatory signals. Endocrinology, 148(11), 5230–5237. doi:10.1210/en.2007-0394

Yamamoto, M., Sato, S., Hemmi, H., Hoshino, K., Kaisho, T., Sanjo, H., Takeuchi, O., Sugiyama, M., Okabe, M., Takeda, K., & Akira, S. (2003). Role of adaptor TRIF in the MyD88-independent toll-like receptor signaling pathway. Science, 301(5633), 640–643. doi:10.1126/science.1087262

Yamawaki, Y., Kimura, H., Hosoi, T., & Ozawa, K. (2010). MyD88 plays a key role in LPS-induced Stat3 activation in the hypothalamus. American Journal of Physiology - Regulatory, Integrative and Comparative Physiology, 298(2), R403.

Zhang, Y., Chen, K., Sloan, S. A., Bennett, M. L., Scholze, A. R., O’Keeffe, S., Phatnani, H. P., Guarnieri, P., Caneda, C., Ruderisch, N., Deng, S., Liddelow, S. A., Zhang, C., Daneman, R., Maniatis, T., Barres, B. A., & Wu, J. Q. (2014). An RNA-Sequencing Transcriptome and Splicing Database of Glia, Neurons, and Vascular Cells of the Cerebral Cortex. The Journal of Neuroscience, 34(36), 11929–11947. doi:10.1523/JNEUROSCI.1860-14.2014

Zhu, X., Levasseur, P. R., Michaelis, K. A., Burfeind, K. G., & Marks, D. L. (2016). A distinct brain pathway links viral RNA exposure to sickness behavior. Sci Rep, 6, 29885. doi:10.1038/srep29885

